# Estimation of neural network model parameters from local field potentials (LFPs)

**DOI:** 10.1101/564765

**Authors:** Jan-Eirik W. Skaar, Alexander J. Stasik, Espen Hagen, Torbjørn V. Ness, Gaute T. Einevoll

## Abstract

Most modeling in systems neuroscience has been *descriptive* where neural representations, that is, ‘receptive fields’, have been found by statistically correlating neural activity to sensory input. In the traditional physics approach to modelling, hypotheses are represented by *mechanistic* models based on the underlying building blocks of the system, and candidate models are validated by comparing with experiments. Until now validation of mechanistic cortical network models has been based on comparison with neuronal spikes, found from the high-frequency part of extracellular electrical potentials. In this computational study we investigated to what extent the low-frequency part of the signal, the local field potential (LFP), can be used to infer properties of the neuronal network. In particular, we asked the question whether the LFP can be used to accurately estimate synaptic connection weights in the underlying network. We considered the thoroughly analysed Brunel network comprising an excitatory and an inhibitory population of recurrently connected integrate-and-fire (LIF) neurons. This model exhibits a high diversity of spiking network dynamics depending on the values of only three synaptic weight parameters. The LFP generated by the network was computed using a hybrid scheme where spikes computed from the point-neuron network were replayed on biophysically detailed multicompartmental neurons. We assessed how accurately the three model parameters could be estimated from power spectra of stationary ‘background’ LFP signals by application of convolutional neural nets (CNNs). All network parameters could be very accurately estimated, suggesting that LFPs indeed can be used for network model validation.

**Significance statement:** Most of what we have learned about brain networks *in vivo* have come from the measurement of spikes (action potentials) recorded by extracellular electrodes. The low-frequency part of these signals, the local field potential (LFP), contains unique information about how dendrites in neuronal populations integrate synaptic inputs, but has so far played a lesser role. To investigate whether the LFP can be used to validate network models, we computed LFP signals for a recurrent network model (the Brunel network) for which the ground-truth parameters are known. By application of convolutional neural nets (CNNs) we found that the synaptic weights indeed could be accurately estimated from ‘background’ LFP signals, suggesting a future key role for LFP in development of network models.

## 1 Introduction

The traditional physics approach to modeling typically involves four steps: (i) A hypothesis is formulated in terms of a candidate *mechanistic* mathematical model, that is, a model based on interactions between building blocks of the system, (ii) predictions of experimentally measurable quantities are calculated from the model, (iii) the predictions are compared with experiments, and (iv) if necessary, the hypothesis is adjusted, that is, a new candidate model is proposed. In neuroscience, a *descriptive* or *statistical* approach has been more common, in particular in systems neuroscience aiming to understand neural network behaviour *in vivo*. Here statistical techniques are used to look, for example, for correlations between measured neural activity and sensory stimuli presented to the animal to estimate receptive fields (Dayan and Abbott, 2001, Ch. 2). While descriptive models can inform us about neural representations in various brain areas, they do not as mechanistic models inform about the biological mechanisms underlying these representations.

At present, mechanistic network models mimicking specific neural circuits are scarce. For small networks like the circuit in the crustacean stomatogastric nervous system comprising a few tens of neurons, some excellent models have been developed (Marder and Goaillard, 2006). For cortical networks important pioneering efforts to construct comprehensive networks with tens of thousands of neurons mimicking cortical columns in mammalian sensory cortices, have been pursued, e.g., Traub et al. (2005); Potjans and Diesmann (2014); Markram et al. (2015); Arkhipov et al. (2018). These models were found to predict spiking activity in rough qualitative accordance with some observed population phenomena (spiking statistics, spike oscillations, …). Fitting of cortical network models to trial-averaged multi-unit activity (MUA) recorded in somatosensory cortex has been pursued for population firing-rate models (Blomquist et al., 2009). However, we do not yet have validated, general-purpose network models that accurately predict experimentally recorded neural activity both in the various ‘background’ states and as a response to sensory stimulation.

The cortical models above have been compared with experimental spiking activity, that is, the high-frequency part of extracellular electrical potentials. The low-frequency part, the local field potential (LFP), in contrast largely reflects how synaptic inputs are processed by dendrites in the populations of neurons surrounding the electrode contacts (Buzsáki et al., 2012; Einevoll et al., 2013; Pesaran et al., 2018). Several methods for analysis of cortical LFP signals have been developed, see Einevoll et al. (2013); Pesaran et al. (2018) for reviews. However, the LFP signal has only rarely been used to validate specific mechanistic models for cortical networks, but see Mazzoni et al. (2008, 2011).

In the present work we explore to what extent the LFP signal generated by a neuronal network model can be used to extract the connectivity parameters of the same network. As a model network we consider the so-called Brunel network comprising an excitatory and an inhibitory population of recurrently connected integrate-and-fire (LIF) neurons (Brunel, 2000). Point neurons do not generate extracellular potentials, however, and to compute corresponding LFPs we use a hybrid LFP scheme (Hagen et al., 2016): First the spiking activity is computed by use of the simulator NEST (Kunkel et al., 2017), and next the computed spikes are replayed as presynaptic spikes onto biophysically detailed multicompartmental neuron models to compute the LFP using LFPy (Lindén et al., 2014; Hagen et al., 2018). The LFP generated by a network depends crucially on the level of temporal correlations of synaptic input onto the neurons (Lindén et al., 2011; Łęski et al., 2013; Mazzoni et al., 2015; Hagen et al., 2016). Thus the LFPs generated by the Brunel network will, as the spiking activity, vary strongly between the different network states as obtained for different choices of network model parameters.

We assess how well network model parameters can be estimated from the stationary ‘background’ LFP signal. For this, we first train *convolutional neural nets* (CNNs) (Rawat and Wang, 2017) with LFP training data for which the underlying model parameters are known, and then test the accuracy of parameter estimation on a separate set of LFP test data. As it turns out, a relatively simple CNN is sufficient for the task and is indeed found to accurately estimate the network model parameters. Thus for the present example, the LFP signal contains sufficient information to accurately recover the underlying model parameters. This suggest that not only spiking data, but also LFPs, can be used to validate candidate network models.

## 2 Methods

### 2.1 Point-neuron network model

The Brunel network (Brunel, 2000) consists of two local populations, one with excitatory and one with inhibitory neurons. These populations of size *N*_E_ and *N*_I_, respectively, consist of leaky integrate-and-fire (LIF) neurons interconnected with current-based delta-shaped synapses. Inputs from external connections are modeled as uncorrelated excitatory synaptic input currents with activation governed by a fixed-rate Poisson process with rate *ν*_ext_.

The sub-threshold dynamics of the point-neurons obey a first-order differential equation, cf. Equation (1) and 2 in Table 1. When the membrane potential of a neuron reaches its firing threshold *θ*, the neuron emits a spike, the synapses onto all its postsynaptic neurons are activated after a time delay *t*_d_, and the neuron’s membrane potential is clamped to a potential *V*_reset_ for a refractory period of *t*_ref_. Each neuron receives a fixed number of incoming connections (fixed in-degree) from a fraction *ϵ* of all other local neurons in the network in addition to the external input. The synaptic connection strengths are constant for each population, for excitatory neurons and external input it is given by *J*_E_ = *J* and for inhibitory neurons *J*_I_ = *−gJ*. The amount of input the local neurons receive from the external population is determined by the parameter *η* = *ν*_ext_*/ν*_thr_, where *ν*_thr_ = *θ/*(*J τ*_m_) is the minimum constant rate input that by itself will drive a neuron to its firing threshold, and *τ*_m_ is the membrane time constant. A complete description of the point-network model is given in Table 1, with specific parameter values given in Table 2.

**Table 1:**
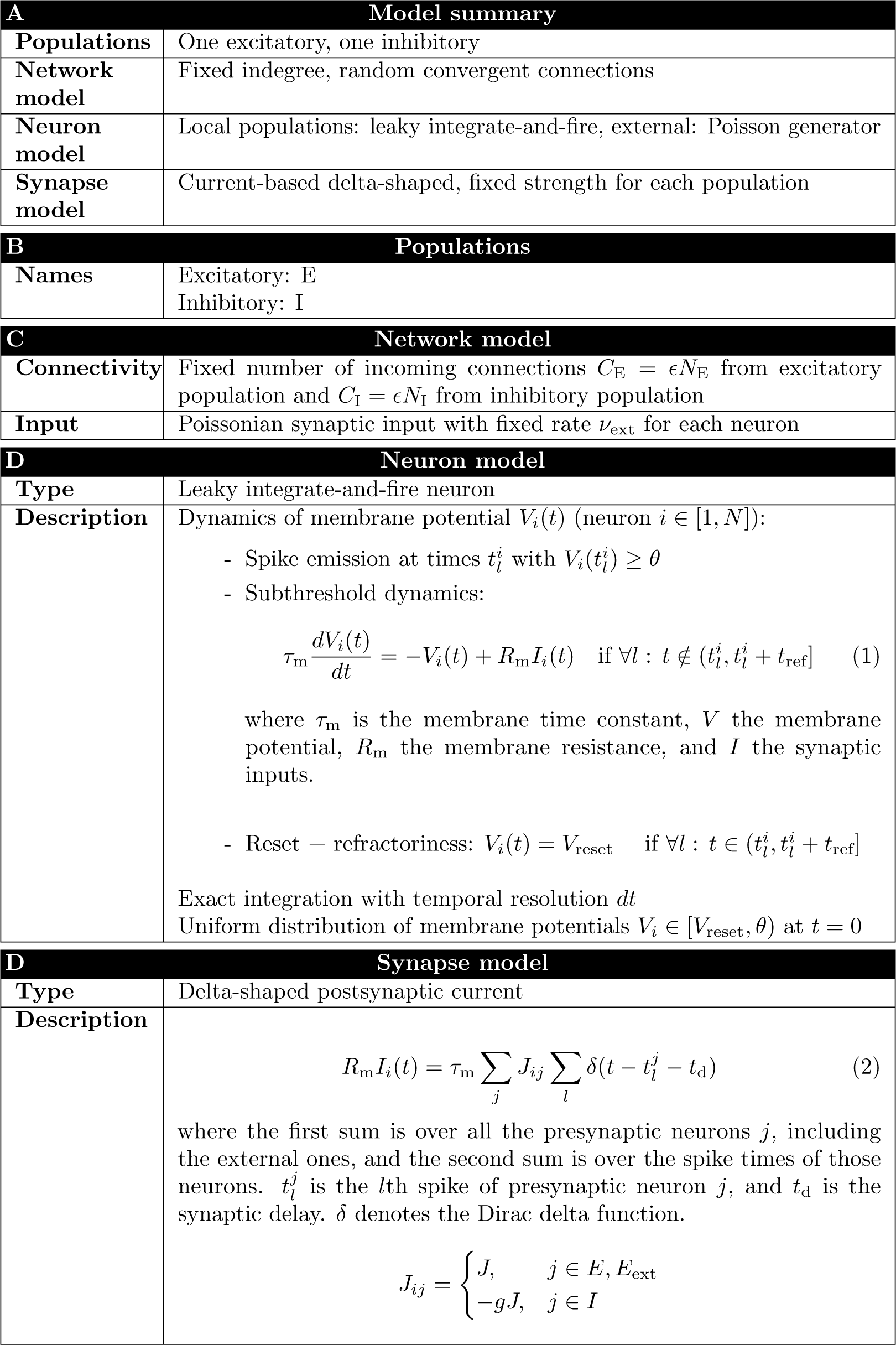
Description of point-neuron network following the guidelines of Nordlie et al. (2009).

**Table 2:** Point-neuron network parameters.

| **Point-neuron parameters** |  |  |
| --- | --- | --- |
| Symbol | Description | Value |
| $\eta$ | relative amount of external input | [0.8, 4.0] |
| $g$ | relative strength of inhibitory synapses | [3.5, 8.0] |
| $J$ | absolute excitatory strength | [0.05, 0.4] mV |
| $\tau_m$ | membrane time constant | 20 ms |
| $C_m$ | membrane capacitance | 250 pF |
| $t_d$ | synaptic delay period | 1.5 ms |
| $t_{ref}$ | absolute refractory period | 2 ms |
| $\theta$ | firing threshold | 20 mV |
| $V_{reset}$ | reset membrane potential | 10 mV |
| $N_E$ | number of excitatory neurons | 10000 |
| $N_I$ | number of inhibitory neurons | 2500 |
| $\epsilon$ | connection probability | 0.1 |
| $C_E$ | number of incoming excitatory synapses | 1000 |
| $C_I$ | number of incoming inhibitory synapses | 250 |
| **Simulation parameters** |  |  |
| Training and validation data |  |  |
| $T_{sim}$ | simulation duration | 3 s |
| $T_{transient}$ | start-up transient duration | 150 ms |
| $dt$ | time resolution | 0.1 ms |
| Model exploration data |  |  |
| $T_{sim}$ | simulation duration | 30.5 s |
| $T_{transient}$ | start-up transient duration | 500 ms |
| $dt$ | time resolution | 0.1 ms |

### 2.2 Forward-model predictions of LFPs

In order to compute local field potentials (LFPs) from the point-neuron network, we utilized the recently introduced ‘hybrid LFP scheme’ (Hagen et al., 2016) (github.com/INM-6/hybridLFPy), illustrated in Figure 1. The scheme allows for the decoupling of the simulation of spiking dynamics (here computed using point neurons) and predictions of extracellularly recorded LFPs. The latter part relies on reconstructed cell morphologies and multicompartment modeling in conjunction with an electrostatic forward model. As the complete description of the scheme (including the biophysics-based forward model) and its application with a cortical microcircuit model (Potjans and Diesmann, 2014) is given in Hagen et al. (2016), we here only briefly summarize the main steps taken to predict LFPs from the two-population network described above: To represent each network population we chose one layer-4 pyramidal neuron and one interneuron reconstruction for the excitatory and inhibitory populations, respectively (Figure 1B). The corresponding morphology files L4E_53rpy1_cut.hoc and L4I_oi26rbc1.hoc were also used in Hagen et al. (2016) (cf. their Table 7), but the apical dendrite of the pyramidal neuron was cut to make it shorter to better fit our smaller column. The somatic positions of all *N*_E_ + *N*_I_ neurons were drawn randomly with homogeneous probability within a cylinder with radius *r* and height ∆*z* (Figure 1B). Each excitatory cell morphology was oriented with their apical dendrite pointing upwards in the direction of the positive *z−*axis and rotated with a random angle around that axis, while inhibitory neurons were rotated randomly around all three axes. The membranes of each morphology were fully passive, with the same membrane time constant *τ*_m_ as in the point-neuron network.

**Figure 1:**
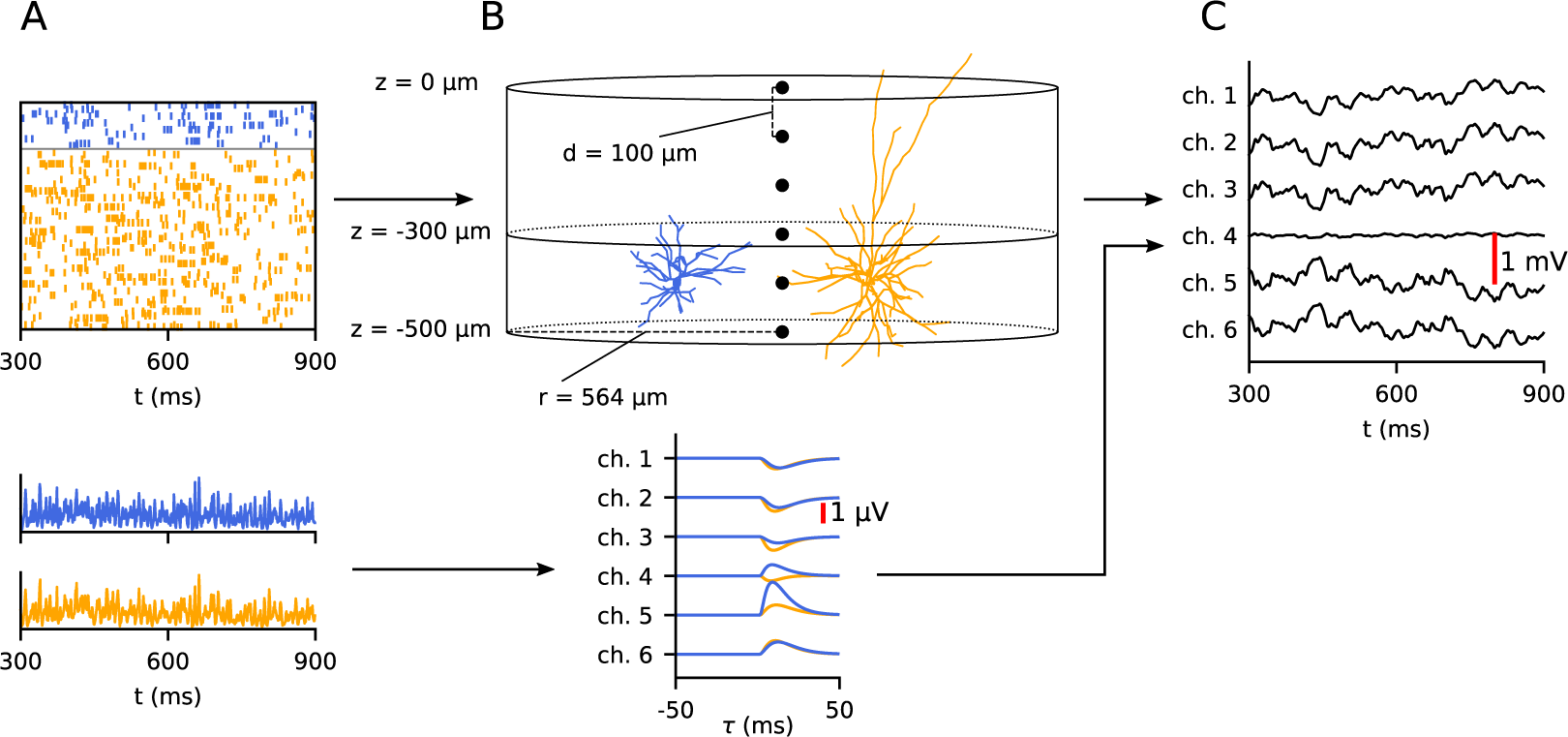
Overview of hybrid scheme for computing local field potentials (LFPs). Top row: First, the dynamics of a network is simulated using a point-neuron simulation (**A**), and the resulting spike times are saved to file. Orange and blue color indicate excitatory and inhibitory neurons. In a separate simulation, the obtained spike times are replayed as synaptic input currents onto reconstructed neuron morphologies representing postsynaptic target neurons (**B**, only one excitatory in orange and one inhibitory neuron in blue are shown). Based on the resulting transmembrane currents of the postsynaptic target neurons in this second simulation, the LFP is calculated **(C)**. Bottom row: Prediction of LFPs from population firing histograms. Instead of running the full hybrid scheme, the LFP can be predicted by the convolution of the population firing histograms (lower figure in **A**) with kernels representing the average contribution to the LFP by a single spike in each population (lower figure in **B**). These kernels are computed using the hybrid scheme, see Hagen et al. (2016, Figure 13).

In the present hybrid scheme the activity in the LFP-generating populations of multicompartment neurons are obtained by mapping spikes generated by individual LIF neurons in the point-neuron network to synapse activation times at specific positions on their equivalent multicompartment neurons. To obtain the synaptic connectivity onto the different positions on the morphologies of the multicompartment neurons, we defined an ‘upper’ and ‘lower’ layer (homologous to e.g., layer 2/3 and 4) on the depth intervals [0*, z*_1_) and [*z*_1_*, z*_2_), see Figure 1B. The layer-specificity of connections (Hagen et al., 2016, p. 4470–4473) was equal between layers for excitatory synapses onto excitatory cells, otherwise all other synapses were made in the lower layer. Within each layer, the probabilities for synaptic connections were proportional to the surface area of each compartment normalized by the total compartment surface area within the layer. Only inhibitory synapses were allowed on the soma compartments. The per-neuron synaptic in-degrees were preserved from the network. As the delta-shaped postsynaptic currents (PSCs) of the point-neuron network cannot be accurately represented in the multicompartment neuron modeling scheme due to numerical discretization of time, alpha-function shaped PSCs (Equation (7) in Table 3) with synaptic time constant *τ*_s_ were used instead. The amplitude of the PSCs was chosen so that the total transferred charge is equal for both synapse types (thus preserving the total synaptic input current between the network and multicompartment neurons). A full description of the multi-compartment neuron model is given in Tables 3 and 4.

**Table 3:** Description of multi-compartment neuron populations.

| A Model summary |  |
| --- | --- |
| Populations | Local excitatory and inhibitory populations |
| Neuron model | Multi-compartment neurons with passive cable formalism |
| Synapse model | Current-based α-function shaped, fixed strength for each population |
| Topology | Cylinder of 1 mm<sup>2</sup> cross-section with somas of both populations positioned in single layer of thickness 0.1 mm. |
| B Neuron models |  |
| Type | Reconstructed multi-compartment morphologies with passive electrical properties |
| Description | For each neuron, the membrane potential $V_n$ of compartment $n$ connected to $m$ other compartments $k$, with a surface area $a_n$, length $l_n$ and diameter $d_n$ is given by:<br>$\sum_{k=1}^m g_{akn}(V_k - V_n) = C_{mn} \frac{dV_n}{dt} + I_{mn} \quad (3)$ $C_{mn} = c_m a_n \quad (4)$ $g_{akn} = \pi(d_n^2 + d_k^2)/(4r_a(l_n + l_k)) \quad (5)$ $I_{mn} = g_{Ln}(V_n - E_L) + \sum_j I_{jn} , \quad (6)$<br>where for compartment $n$, $C_{mn}$ is the membrane capacitance, $g_{akn}$ the axial conductance from compartment $k$, $I_{mn}$ the membrane current, $g_{Ln}$ the membrane leak conductance, $E_L$ the extracellular reversal potential, and $I_{jn}$ the synaptic current from presynaptic neuron $j$. |
| C Synapse model |  |
| Synapse type | α-function shaped postsynaptic current |
| Description | $I(t) = H(t - t_a)JCte^{1-t/\tau_s} \quad (7)$ $H(t) = 0 \text{ for } t \leq 0, \text{ otherwise } 1 . \quad (8)$<br>Here $t_a$ is the activation time of the synapse, $J$ the synaptic strength, and $\tau_s$ is the synaptic time constant. $C$ is a constant chosen so that $JC \int_0^\infty te^{1-t/\tau_s} dt = C_m J$, assuring that the same total charge is transferred as in the δ-function synapse in the point-neuron network. |
| D Topology |  |
| Type | Cylinder with radius $1/\sqrt{\pi}$ mm and height 0.5 mm containing two vertical sections |
| Description | - Cylinder extends from $z=-500 \mu\text{m}$ to $z = 0$<br>- All somas are randomly placed with a uniform distribution within the boundaries $r \leq 564 \mu\text{m}$ and $-450 \mu\text{m} \leq z \leq -350 \mu\text{m}$<br>- Two regions separated by the plane $z = -300 \mu\text{m}$<br>- Synapses on inhibitory neurons are placed in lower region<br>- Inhibitory synapses on excitatory neurons are placed in lower region<br>- Excitatory synapses on excitatory neurons are split equally between regions |

**Table 4:** Multi-compartment neuron parameters.

| Multi-compartment neuron parameters |  |  |
| --- | --- | --- |
| Symbol | Description | Value |
| $\tau_m$ | membrane time constant | 20 ms |
| $C_m$ | membrane capacitance | 1.0 $\mu\text{F}/\text{cm}^2$ |
| $R_m$ | membrane resistivity | $\tau_m/C_m$ |
| $R_a$ | axial resistivity | 150 $\Omega\text{cm}$ |
| $\tau_s$ | synaptic time constant | 5 ms |
| $E_L$ | passive leak reversal potential | 0 mV |
| $V_{\text{init}}$ | membrane potentials at $t = 0$ ms | 0 mV |
| $\sigma_e$ | extracellular conductivity | 0.3 $\text{Sm}^{-1}$ |

The presently used choice of current-based synapses and morphologies with passive membranes in the multicompartment neuron models introduces a linear relationship between any presynaptic spike event and contributions to the LFP resulting from evoked currents in all postsynaptic multicompartment neurons. Thus the LFP contribution 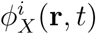 at position **r** from a single presynaptic point-neuron neuron *i* in population *X* can, in general, be calculated by the convolution of its spike train 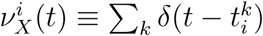 with a unique kernel 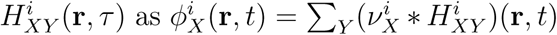. This kernel encompasses effects of the postsynaptic neuron morphologies and biophysics, the electrostatic forward model, the synaptic connectivity pattern, conduction delay and PSCs.

The resulting LFP due to spikes in a presynaptic population *X* is then given by (Hagen et al., 2016)

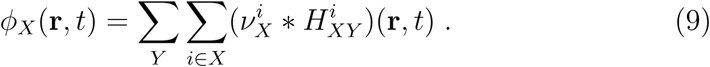

The evaluation of this sum is computationally expensive for large population sizes. For our purposes where the calculation of of LFP signals lasting seconds must be repeated tens of thousands of times to have training and test data for the CNNs, this scheme is not feasible.

Following Hagen et al. (2016, Figure 13) we instead use a firing-rate approximation and compute the LFP by a convolution of population firing rates 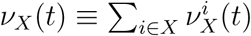 and averaged kernels 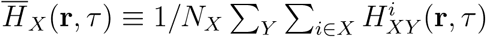
that is,

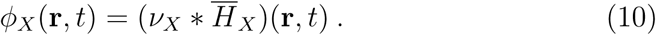

Similar to Hagen et al. (2016), these averaged kernels 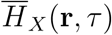 were here measured using the full hybrid-scheme set up by replacing ongoing spiking activity in the point-neuron network populations by fully synchronous spike events, that is, 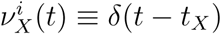 where *t*_*X*_ is the timing of the synchronous event in population *X*. In this way the computational resources needed to run LFP simulations are reduced by several orders of magnitude compared to direct use of Equation 9. To test the accuracy of the approximation of using Equation 10 instead of Equation 9, we compared their LFP predictions for a set of example parameter sets and found in general excellent agreement between the resulting power spectra. A comparison is shown in the lower panels of Figure 5 in Results.

The kernel 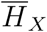 will scale linearly with the postsynaptic strengths of population *X*, and is therefore dependent on the parameters *J* for *X ∈* [*E, I*] and *g* for *X ∈* [*I*]. The kernels were thus computed only once for a set of reference values for *J* and *g*, and for each simulation these reference kernels were scaled accordingly to the particular values of *J* and *g*. The LFP was computed across depth through the center of the cylindric volume with a spatial resolution *d* as illustrated in Figure 1B for the same duration as the network simulations.

### 2.3 Statistical methods

#### LFP spectral analysis

The power spectral densities *P*_*φ*_(**r**, *f*) of LFPs *φ*(**r**, *t*) in each location **r** were estimated using Welch’s average periodogram method (Welch, 1967). For this we used the implementation from the Python SciPy package (Jones et al., 2001–) (`scipy.signal.welch`), with parameters listed in Table 5.

**Table 5:** Parameters for Welch’s method for computing power spectral density (PSD) of LFP.

| Power spectrum estimation |  |  |
| --- | --- | --- |
| Symbol | Description | Value |
| NFFT | window length | 300 ms |
| noverlap | segment overlap | 150 ms |
| Fs | sampling frequency | 1 kHz |
| window | window function | Hann |

##### Statistical measures of activity

Two statistical measures were employed to probe the spiking network activity in the different regions of the parameter space. Simulations of 30.5 seconds of the activity were run and used to calculate the statistics, where the first 500 ms of the simulations were discarded.

The mean network firing rate, including both the excitatory and inhibitory populations was calculated as

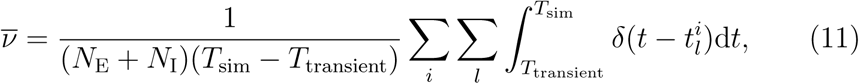

over all neurons *i* and their spikes *l* at spike times 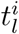. The coefficient of variation (CV) of the inter-spike intervals (ISI) of individual neurons was used as a measure of the irregularity of firing (Grün and Rotter, 2010). The presently used mean CV was defined as

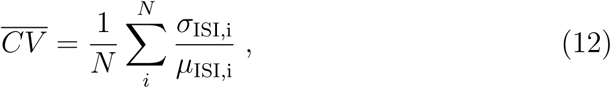

averaged over all neurons *i*.

As a measure of the degree to which the LFP power spectrum is spread out over different frequencies, we employed the entropy of the normalized power spectrum of the LFP measured in the uppermost channel, defined as

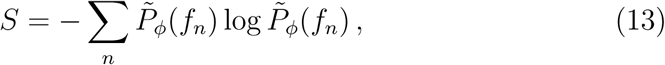

where 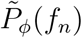 is the power spectrum of the LFP *φ*(**r***, t*) at frequency *f*_*n*_ normalized to unity. Since the power spectrum is computed numerically using Welch’s method, this introduces a discretisation in frequency space.

### 2.4 Simulation of training and validation data

Two different sets of training and validation data were created for this study. The first set was generated by a wide parameter space wherein the pointneuron network parameters *η* ∈ [0.8, 4.0], *g* ∈ [3.5, 8.0] and *J* ∈ [0.05, 0.4]mV were 50000 parameter triplets were randomly selected with homogeneous probability. This first set thus encompassed the four different activity states that are displayed by the Brunel network, illustrated in Figure 2. These activities include synchronous regular (SR), asynchronous irregular (AI), synchronous irregular (SI) with either slow or fast oscillations. This wide-spanning parameter space is illustrated by the orange outline. The second training and validation data set was generated by drawing 50000 parameter combinations from a narrower parameter space where *η* ∈ [1.5, 3.0], *g* ∈ [4.5, 6.0] and *J* ∈ [0.1, 0.25] mV. This second data set encompassed AI activity states, as illustrated by the blue outline in Figure 2. All other parameters (Table 2) were kept constant in the simulations, which each was run for a duration of *T*_sim_ = 3 s. Start-up transients with duration *T*_transient_ = 150 ms were discarded. LFP signals for all spiking output were computed as outlined above, and as final training and validation data we estimated the power spectrum *P*_*φ*_(**r**, *f*) in each LFP channel.

**Figure 2:**
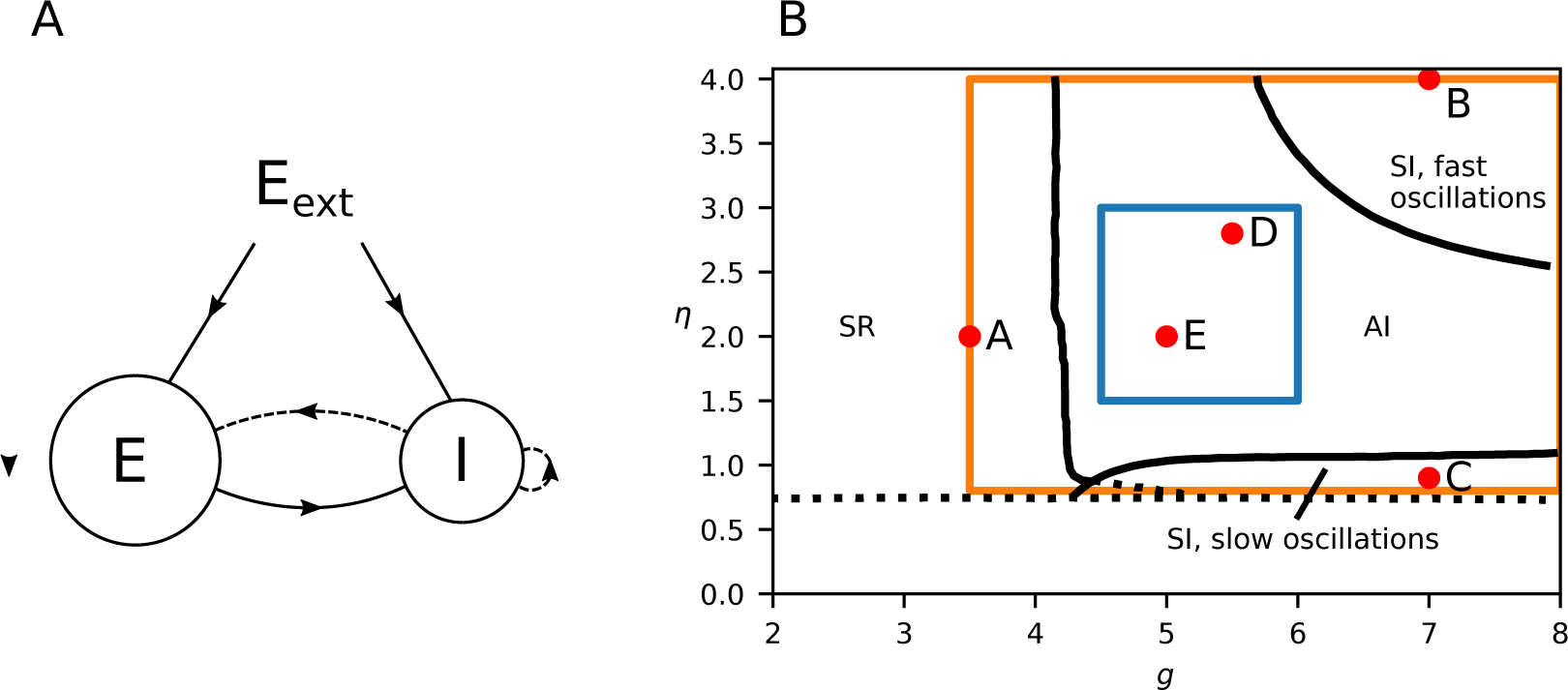
Brunel model network and phase diagram. **A**, Illustration of network. Solid lines represent excitatory connections, dashed lines inhibitory connections. **B**, Phase diagram, adapted from Brunel (2000), Figure 2A. Different network states arise depending on the parameters *η* = *ν*_ext_/*ν*_thr_ and *g* (where in the present example a fixed synaptic delay *t*_d_ of 1.5 ms is used). SR stands for synchronous regular, SI for synchronous irregular, and AI asynchronous irregular. Orange box shows the extent of of parameters we simulated and blue box when we restricted the simulations to the AI state. Note that this plot shows a slice of the parameter space for a given value of *J* = 0.1. We considered different values of *J* in the study, so the actual parameter space is a cube, with the third axis being in the *J*-direction. The red dots labeled A–E indicate the *η* and *g* values of the example activities shown in Figure 5.

### 2.5 Parameter estimation by convolutional neural networks

The CNN architecture is illustrated in Figure 3 and fully described in Table 6, and was set up using the Keras machine learning framework (Chollet et al., 2015) running on top of TensorFlow (Abadi et al., 2015). It consisted of three convolutional layers with 20 filters, each followed by max pooling layers, and two fully connected layers before the output layer. The rectified linear unit (ReLU) function *f* (*x*) = max(0*, x*) was used as the activation function for all layers apart from the output layer, and biases were only used in the fully connected layers. As input, it took the PSD of each LFP channel, a 6 by 151 matrix. The convolutions were done in one dimension, with kernels extending over all LFP channels. There were two fully connected layers, with 128 nodes each, before the output layer consisting of 3 nodes. Each node in the output layer corresponded to a single parameter *η*, *g* and *J*.

**Table 6:** **Detailed specification of presently used convolutional neural network (CNN).** The convolutional kernel dimensions are given as [frequency, channels in, channels out], the strides and window sizes are given in the frequency dimension.

| Convolutional neural network |  |
| --- | --- |
| Layer | Description |
| Conv. layer 1 | kernel size: 12x6x20<br>stride: 1<br>activation: ReLU<br>bias: no |
| Max pool 1 | window size: 2<br>stride: 2 |
| Conv. layer 2 | kernel size: 3x20x20<br>stride: 1<br>activation: ReLU<br>bias: no |
| Max pool 2 | window size: 2<br>stride: 2 |
| Conv. layer 3 | kernel size: 3x20x20<br>stride: 1<br>activation: ReLU<br>bias: no |
| Max pool 3 | window size: 2<br>stride: 2 |
| Dense layer 1 | nodes: 128<br>activation: ReLU<br>bias: yes |
| Dense layer 2 | nodes: 128<br>activation: ReLU<br>bias: yes |
| Output layer | nodes: 3<br>activation: None<br>bias: no |

**Figure 3:**
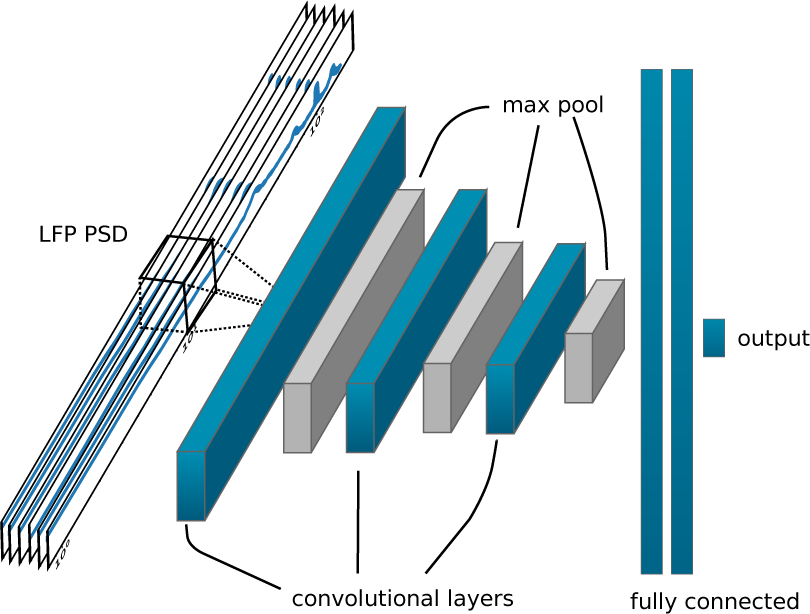
Illustration of convolutional neural network (CNN). The PSDs of all six LFP channels are taken as input. The three convolutional layers consist of 20 filters each, and are followed by max pooling. Two fully connected layers precede the output layer which consists of 3 nodes, one for each parameter.

The LFP PSD was normalized for each channel by the mean of the sum of the PSD over all frequencies, serving to diminish the variation in amplitude across the different LFP PSDs input to the network, while keeping the variation in amplitude across channels for each single LFP PSDs. For labels, each parameter was linearly mapped to the interval [0, 1].

The network was trained by batch gradient descent on 40000 of the simulated LFPs, while the final 10000 simulated LFPs were reserved for validation. To train the CNN, we required a *loss function* which was minimized during training. We defined the *loss* as the mean squared error of the estimator

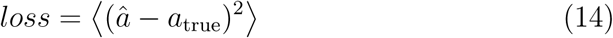

where â is the estimate (output from the CNN) and *a*_true_ is the truth (‘ground-truth’ value) of any network parameter *a*.

The Adam optimizer (Kingma and Ba, 2014) was used, with a batch size of 100, learning rate of 0.001 and the *β*_1_, *β*_2_ and *E*_Adam_ parameters at their suggested default values. The networks were trained for 400 epochs, and the network weights with the lowest loss were saved.

### 2.6 Effect of duration of LFP signals

It was *a priori* not known what duration of the data are required to obtain stable results. To test this, the duration of each LFP simulation was successively extended, the PSD of the LFP was computed using the Welch method (cf. Section 2.3), and the CNN was trained with the data to predict the three parameters simultaneously. The test loss during training is shown in Figure 4A. Overall, the loss decreased with training duration and reached a plateau after a certain amount of training epochs. Note that with increasing stimulation duration of the data, the loss got smaller. This was due the larger variation in the computed PSDs for shorter simulations. With longer duration of the LFP signals used in the PSD calculations, the variations will be smaller. The results in the figure suggested that a simulation duration of about 1800 ms would be a good choice, as shorter simulation times decreased the performance. Figure 4B shows the scaling of the minimal test loss (that is, loss obtained in the limit where more training epochs do not improve results) as a function of simulation duration. The 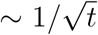 least squares fit was motivated by the scaling of the error of the mean, which gives the square root dependence of the standard error of the mean. This scaling assumes uncorrelated experiments, which is not the case when using Welch’s method as we do. Nevertheless, the fact that this scaling held also for our estimator gave a hint when the uncertainty is still limited by statistical fluctuations.

**Figure 4:**
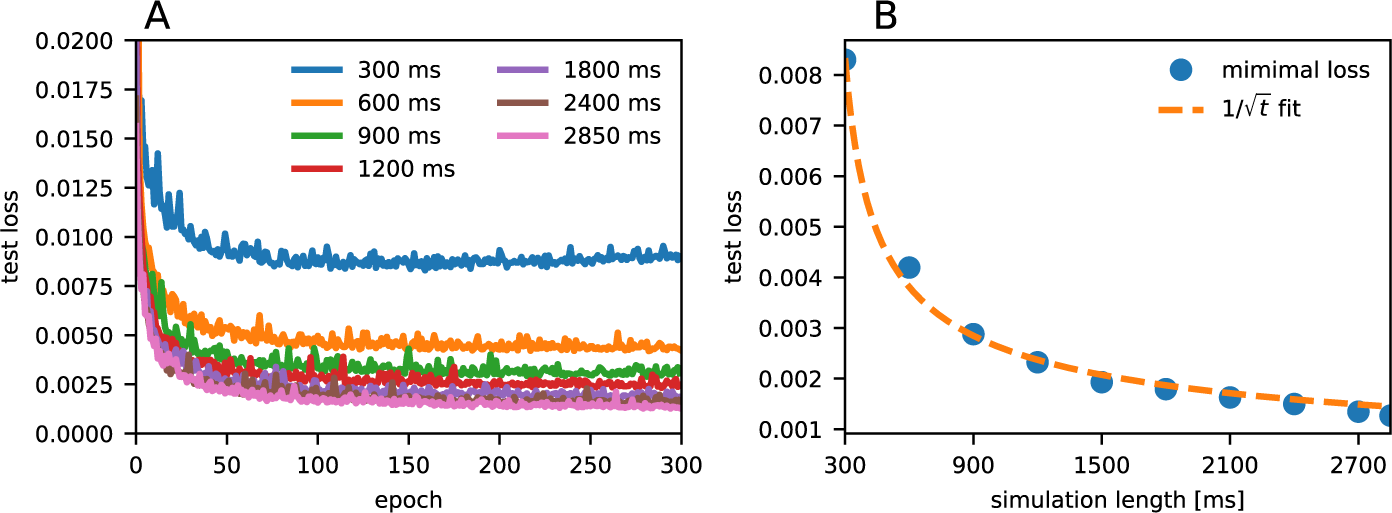
**A**, Test loss as a function of number of training epochs of the CNN for different simulation lengths. **B,** Minimal loss (that is, smallest loss in panel A) as a function of simulation length. A function with 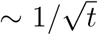 shape was fitted to the data to illustrate that the scaling is dominated by limited statistics. The *R*^2^ score was 0.994.

### 2.7 Technical details

#### 2.7.1 Reproducibility

The simulated results presented here were done using Python v2.7.12. All point-network simulations were done with the NEST simulator v2.12.0 (Kunkel et al., 2017). The forward-modeling of the LFP was done using hybridLFPy v0.1.3 (Hagen et al., 2016), with NEURON v7.5 (Hines et al., 2009). All simulations were run on the Stallo high-performance computing cluster consisting of 2.6 GHz Intel Xeon E5 2670 and 2.8 GHz Intel Xeon E5 2680 CPUs.

The convolutional neural networks were trained using Python v3.5.2 using Keras v2.2 with TensorFlow v1.10.0 as backend.

## 3 Results

The aim of this study is to investigate the possibility of estimating network model parameters for the Brunel two-population spiking-network model (Brunel, 2000) from the stationary ‘background’ LFP signal. We start by describing this spiking model and its salient dynamical properties and further describe how the resulting spikes can be used in a hybrid scheme to calculate associated LFPs (Hagen et al., 2016). Then we discuss the estimation performance of a convolutional neural network (CNN) to predict network parameters of the Brunel network model based on LFP data only.

### 3.1 Network model and LFPs

The presently used Brunel network produces four different network states dependent on the post-synaptic potential amplitude of excitatory connections *J*, the ratio of inhibitory to excitatory connection strength *g*, and the strength of the external input *η* relative to the threshold rate, see Figure 2. In the synchronous regular (SR) state the neurons fire regularly and in synchrony. The asynchronous regular (AR) state is characterised by regularly firing neurons which largely fires unsynchronised with respect to each other. The third state is the asynchronous irregular (AI) state, where individual neurons have an irregular firing rate and very little synchronization. The fourth state is the synchronous irregular (SI) state, characterised by oscillatory population firing rates, yet highly irregular firing of individual neurons. Example spike raster plots and population firing rates for SR, AI and SI states of the network are shown in the top rows of Figure 5. AI states are commonly believed to be realised in most healthy neural networks *in vivo*, often characterized by low average pairwise spike-train correlations (see e.g., Ecker et al. (2010)) and irregular spike trains (see e.g., Mochizuki et al. (2016)).

**Figure 5:**
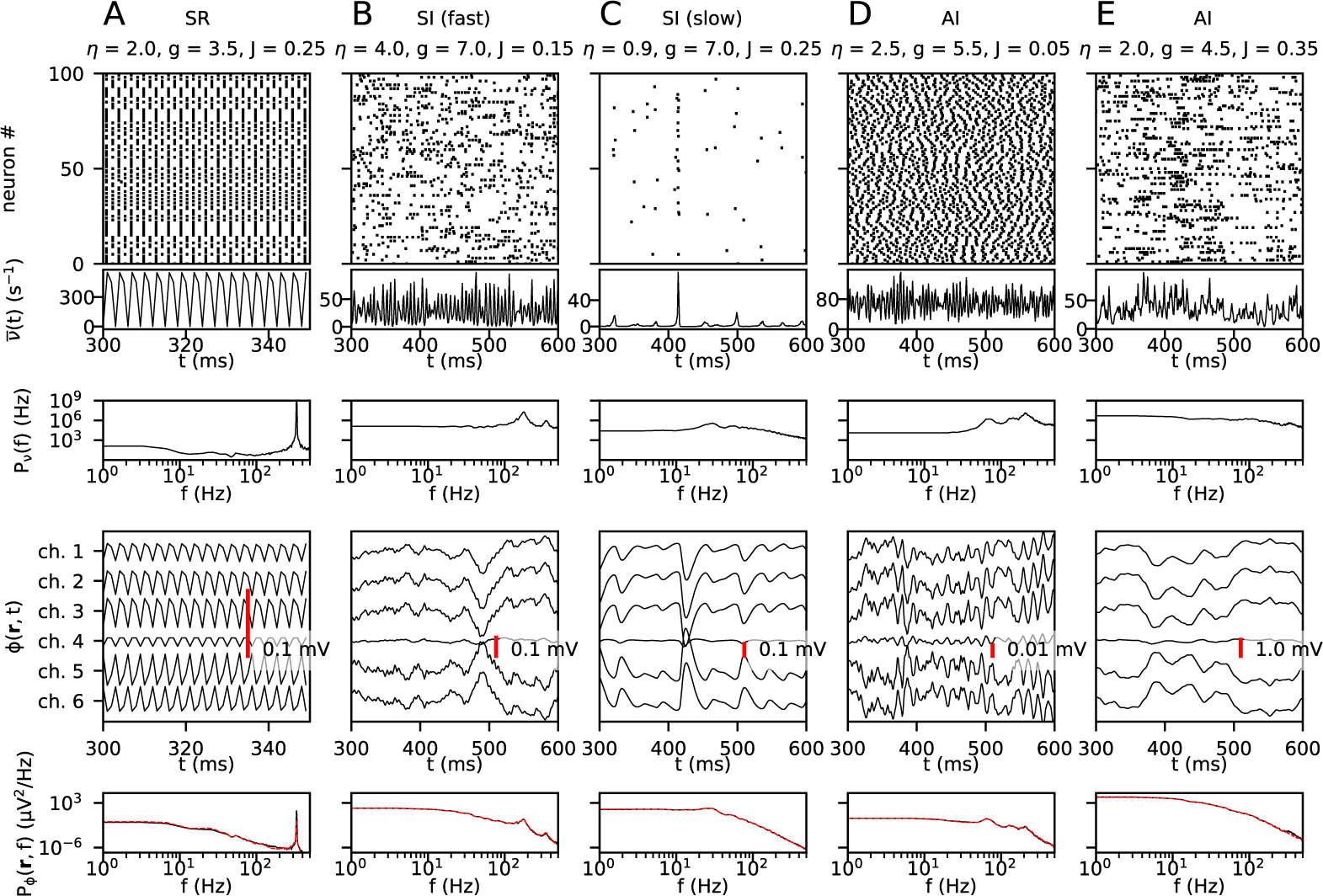
Examples of simulated spiking network activity and LFPs for different sets of network parameters (*η, g* and *J*). For each simulation, **A**–**E**, the first row shows spike trains from 100 randomly selected neurons across both populations. The second and third row show the population firing rate (including both the excitatory and inhibitory neurons) and its power spectral density (PSD). The final two rows show the LFP signal from all six channels and the PSD of channel 1, respectively. The dashed red lines in the lowest panel shows the LFP PSD computed from spikes in individual neurons (Equation 9) rather than with the presently used population firing-rate approach (Equation 10, black lines) which is computationally much less demanding. In general, the agreement is seen to be very high, the only discrepancy is seen for the SR-state example where the height of the peak around 300 Hz differs. The network states for the five examples (SR/SI(fast)/SI(slow)/AI, see text) are indicated at the top.

To generate training and test data for the convolutional neural net (CNN), we simulated the network for many combinations of parameters (*η* ∈ [0.8, 4], *g* ∈ [3.5, 8] and *J* ∈ [0.05, 0.4] mV). This parameter space includes parameter combinations giving three of the states described above: AI, SI, and SR (see orange rectangle in Figure 2). For details of the simulation procedure, see Section 2.4.

The LFPs were simulated using the so-called hybrid scheme introduced by Hagen et al. (2016). In this scheme, neuronal network activity is predicted by a point-neuron network (here the Brunel network), and the corresponding LFPs are estimated in a latter step by ‘playing back’ spike times as activation times of synapses distributed across reconstructed neuron morphologies representative for each population type. The LFP is then computed from the resulting transmembrane currents combined with an electrostatic forward model derived from volume-conductor theory, as detailed in Section 2.2. An overview over the hybrid scheme, including the geometrical organisation of the ‘cortical column’ used in the LFP-generating step, is shown in Figure 1.

#### 3.1.1 Exemplary LFPs for different network states

In the presently used model set-up, the LFP is linearly dependent on the point-neuron network spiking activity (see Section 2.1 and Section 2.2). Any network parameter change affecting ongoing spiking activity will therefore directly affect the LFP. The panels in the lower two rows of Figure 5 show the resulting LFP and LFP power spectra for five different network parameter combinations of *η, g* and *J*.

An example synchronous regular state (SR) is shown in panel A. The simulation showed high regularity and synchrony of the individual spike trains and a strongly oscillating population firing rate. The corresponding LFP generally had a similar time course over all channels, though with opposite phases for the topmost and lower recording channels. The power spectral density (PSD) of the LFP showed a decrease in power with increasing frequency, though with a clear peak at around 333 Hz. This peak was also seen in the PSD of the firing-rate, reflecting the tight relationship between spikes and LFPs.

Two examples of the synchronous irregular state (SI) are illustrated in Figure 5B,C, characterised by synchrony of the firing of neurons while individual neurons fire irregularly. In panel B an example with high firing and fast oscillations is shown. Here the power spectrum of the LFP showed two peaks at around 175 and 350 Hz, respectively. Again, the same peaks are found also in the firing rate spectra. In contrast, panel C shows a low-firing SI state with more slowly varying population firing rates, though without any notable peak in the firing-rate or LFP power spectra.

Two examples of the asynchronous irregular state (AI) are shown in the last two panels (Figure 5D,E). As suggested by the name, this state is defined by lack of synchrony between different neurons and irregular firing patterns of each neuron. For the example in panel D, the firing-rate PSD exhibited three high-frequency peaks, with the peak at the highest frequency (*∼*200 Hz) being highest. The same three peaks were found also in the LFP PSD, but now the peak at the lowest frequency (*∼*70 Hz) was highest. This reflects low-pass filtering effects of the LFP from synaptic and intrinsic dendritic filtering (Lindén et al., 2010; Łęski et al., 2013). In panel E the recurrent excitation *J* is much increased compared to the example in panel D. This combined with a reduction of the relative inhibition *g*, gave much larger LFP signals, as reflected in the high power of the LFP seen for the low frequencies in the LFP PSD. For this parameter set the firing-rate PSD exhibited a broad peak around 100 Hz, but this peak was absent in the corresponding LFP PSD due to synaptic and intrinsic dendritic low-pass filtering.

#### 3.1.2 Model behaviour across parameter space

To extract network model parameters from recordings of neural activity such as the LFP, the network model parameters must necessarily be reflected in these recordings. After the qualitative discussion above, we proceed to discuss how the network behaves over the entire parameter space. We therefore give an overview of how different spike-and LFP-based measures of neural activity vary across parameter space.

##### Spikes

Panel A in Figure 6 shows how the mean network firing rate varied over the parameter space. The overall trend was that with increasing *g*, the firing rate decreased since inhibition was increased. The transition at *g* = 4 resulted from the fact that there are four times more excitatory neurons than inhibitory, and thus for *g* < 4 excitation dominates network behaviour. For *J* ≳ 0.15, three separable regions with smooth transitions emerged: A region of high firing rate on the left border of the plots (*g* ≲ 4), a region of low firing rate on the bottom of the plots (*g* ≳ 5, *η* ≲ 1), and a region of intermediate firing rate in the upper right of the parameter space. For smaller values of *J* (*J* ≲ 0.1) the transition between the high-firing region and the intermediate-firing region became smoother. Thus, large values of *J* amplified the differences between the regions. These distinct regions in the firing-rate phase diagram correspond well with the phase diagrams derived by Brunel (2000), see Figure 2.

**Figure 6:**
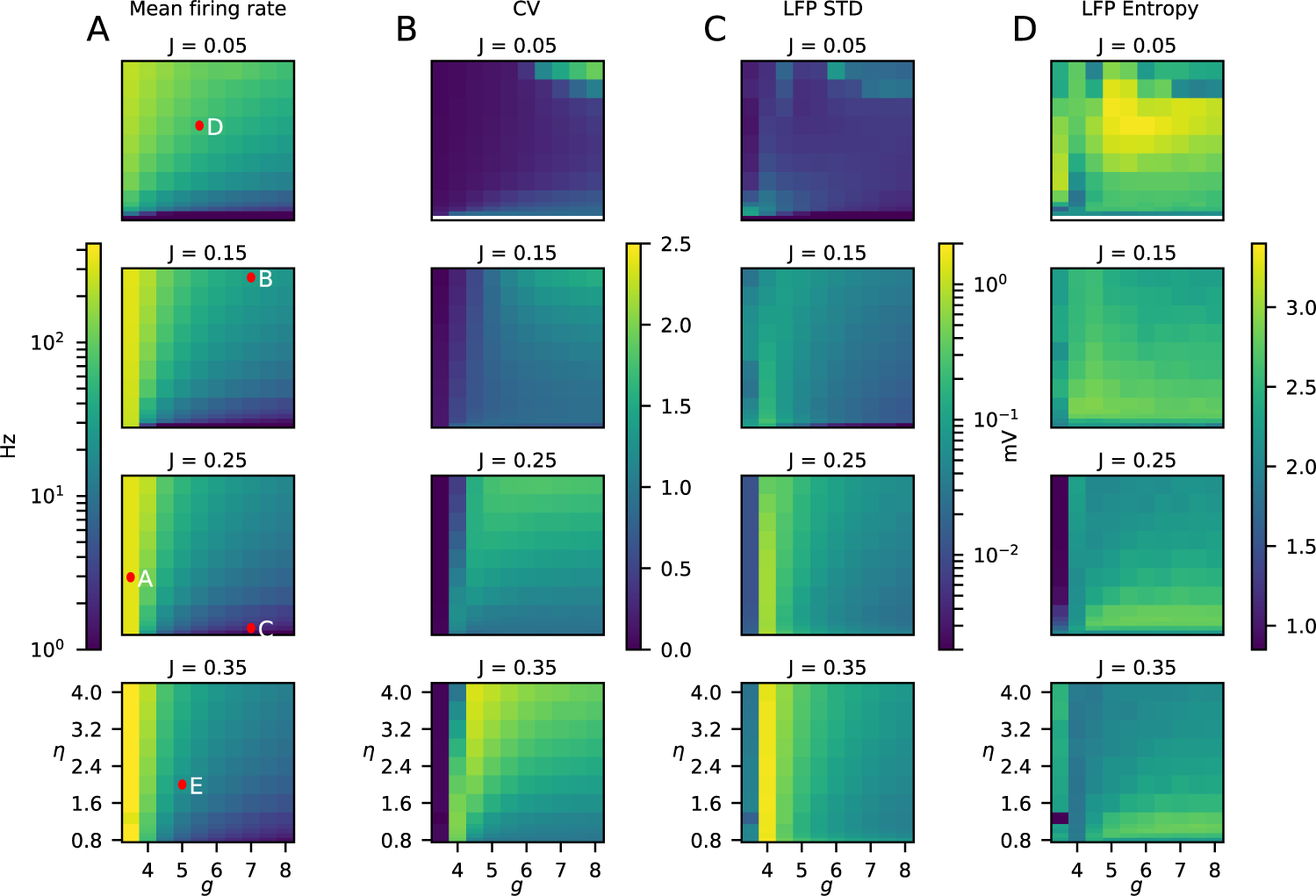
Statistical measures of network activity for different combinations of network parameters (*η, g* and *J*). **A**, Average population firing rates, that is, average firing rate over all neurons and times. The red dots show the parameter values of the examples in Figure 5. **B**, Mean coefficient of variation (Equation (12)) of the inter-spike intervals over all neurons as a measure of the spiking irregularity. **C**, Square root of the variance of the LFP signal integrated over time for the topmost channel (channel 1). This measure corresponds to the square root of integral of the power spectrum of the LFP over all frequencies (Lindén et al., 2011), and is referred to as the standard deviation of the LFP (LFP STD). **D**, LFP Entropy, cf. Equation 13.

Panel B in Figure 6 correspondingly displays the parameter dependence of the average coefficient of variation (CV) of the inter-spike intervals. Similar to the population firing rate, one can see a boundary at about *g ≈* 4 over a large part of the considered parameter range of *J* and *η*. In the region with low *η* and high *g* (*η* ≲ 1.5, *g* ≲ 5) there was also a distinct area with low CV, reflecting the expected lower CV of the slow-oscillation SI state compared to the AI state for larger values of *η*. For small values of *J* (*J* ≲ 0.1), there was a region of larger CV visible in the upper right corner of the parameter space (*g* ≳ 6 and *η* ≳ 3.6). This region overlaps with fast-oscillation SI state described by Brunel (2000) (see phase diagram in Figure 2).

##### LFP

The example LFP patterns in Figure 5 showed substantial variability of the LFPs for different network parameter values. This suggests that it indeed may be possible to estimate network parameters from the LFP. To explore this in more detail, we show in panels C and D of Figure 6 two different measures of LFP signals across the same parameter space.

Panel C shows a measure of the overall signal power of the LFP signal, that is, the standard deviation (STD) for the topmost channel (channel 1). This measure corresponds to the square root of the variance of the LFP signal integrated over all frequencies (Lindén et al., 2011). In panel C a first observation was that large values of the excitatory weight *J* led to higher values of the LFP STD, not surprising given the stronger excitatory synaptic inputs. Likewise, it was seen that the LFP STD generally decreased when inhibition, that is, *g*, increased. Interestingly, despite the very high firing activity for values of *g* smaller than 4, the LFP STD was small for these parameter values. This can be understood by inspection of panel A in Figure 5 which shows results for an example state with *g* = 3.5: Even if there are strong bombardments of synaptic inputs onto the LFP-generating excitatory neurons, the input is so clock-like and regular that there is little power in the LFP signal at the lower frequencies. The only strong LFP signal contribution was obtained for frequencies over *∼*300 Hz, corresponding to the peak seen in the firing rate PSD.

The LFP STD measure considered in panel C measures the overall LFP signal strength across frequencies. In contrast, the measure labeled ‘LFP Entropy’ in panel D measures how much the overall LFP power is spread across the different frequencies, cf. Equation 13 in Methods. The largest entropy value was observed for the smallest excitatory weight (*J* = 0.05 mV), but the detailed parameter dependence of the LFP entropy was not the main point here. The most important observation was that the parameter dependence of LFP Entropy was qualitatively different from the parameter dependence of LFP STD. This implied that the frequency-resolved PSD contained more information regarding the underlying network parameters than either the overall amplitude (LFP STD) or the frequency spread (LFP Entropy) alone. This provided cautious optimism that the variation of the LFP PSD is sufficiently strong across parameter space to allow for estimation of network parameter values with a suitable estimation methods.

### 3.2 Network parameters are accurately estimated from LFP

After this rough survey over how the LFP for the Brunel network model vary across parameter space, we now ask the question: Can the network parameters be estimated from this LFP by use of Convolutional Neural Networks (CNNs)? We chose to use CNNs because they do not rely on manual feature extraction, and our analysis thus do not depend on any assumption of how the model network parameters are reflected in the LFP. Further, we used the power spectral density (PSD) of the LFP for this analysis, that is, used the PSD as input to the CNNs. This approach removes phase information in the LFP. However, since we only considered LFP data from stationary network activity, the hypothesis was that most of the available relevant information regarding network parameters should the contained in the PSD.

Our CNN consisted of three convolutional layers followed by two fully connected layers. An illustration can be seen in Figure 3, and detailed specifications are given in Section 2.5 in Methods. We generated several pairs of training and testing data sets for different scenarios. The parameter space was both sampled randomly and on a regular grid. We also generated a training and test data set on a subset of the parameter space, but with the equal amount of simulations.

While several approaches were tested and compared, we defined the following set-up as the standard set-up: The data was simulated using randomly distributed parameters *η, g* and *J* with a simulated duration of 2.85 seconds for each trial, see Section 2.4. From the simulated LFP, the power spectral densities (PSD) for six recording channels were computed and used as input to the CNN. Then, a single CNN network was trained to predict the parameter vector 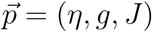 simultaneously, and all three parameters were set to contribute equally to the loss function, Equation 14. To achieve this, the parameter ranges of *η*, *g* and *J* were all scaled to the unit interval [0, 1] for the considered part of the parameter space.

To quantify and illustrate the accuracy of the parameter estimation we used the estimation error â *− a*_true_ where *a*_true_ was the true value and â the estimated value. Figure 7 (orange lines) shows the accuracy of the three network parameters when considering the full parameter space (*η* ∈ [0.8, 4), *g* ∈ [3.5, 8) and *J* ∈ [0.05, 0.4) mV). As observed, the estimation errors are in all cases generally smaller than 5%. Also, the estimations had small biases, that is, the mean errors were close to zero.

**Figure 7:**
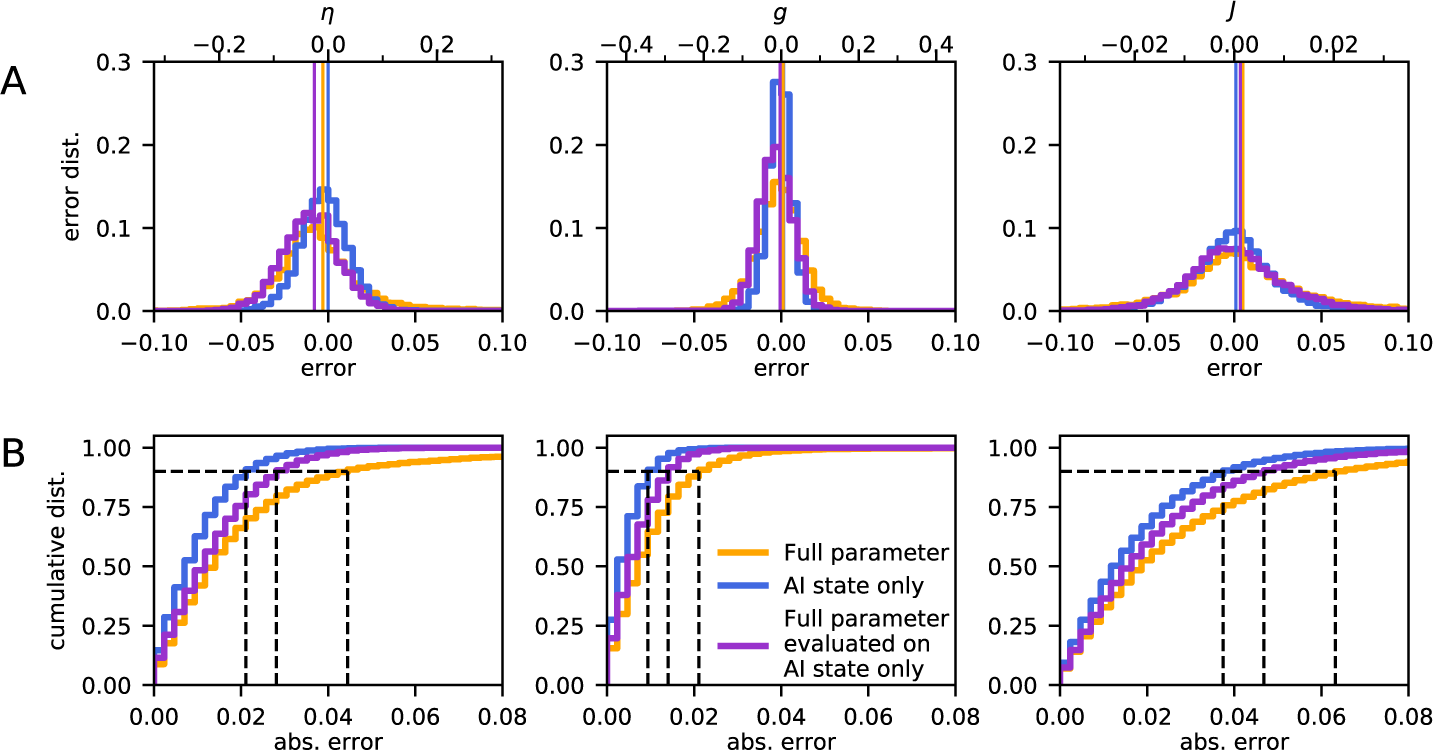
Accuracy of network parameter estimation. **A**, Estimation error distributions for *η, g* and *J* averaged over the entire parameter space. In the plots all parameter ranges were rescaled to the interval [0, 1] for easier comparison on the lower x-axis, the upper x-axis shows the original values The vertical line indicates the mean of both distributions. The orange curve shows the result when using the full parameter set (*η* ∈ [0.8, 4], *g* ∈ [3.5, 8] and *J* ∈ [0.05, 0.4]) and the blue curve when the parameter set only contains the AI state (*η* ∈ [1.5, 3], *g* ∈ [4.5, 6] and *J* ∈ [0.1 − 0.25]). The purple line gives the estimation error of the CNN trained for the full parameter set, but evaluated on the restricted parameter set containing the AI state only. To compare the full parameter data set and the AI-only data set, they were both scaled to the range of the full parameter set. **B**, Cumulative error distributions, the proportion of absolute errors that fall below a given value, also with all parameters rescaled to [0, 1]. The dashed black lines indicate the 90% coverage interval.

The full parameter space considered above covered four of the characteristic network states seen for the Brunel network, see orange rectangle in Figure 2. Here the network-generated LFP can be expected to vary substantially across parameter space making the CNN estimation easier. We thus next explored to what extent CNNs could estimate network parameters within a particular state, that is, the AI state which is thought to be most relevant for cortex.

Training and validation of the CNN were repeated using a second data set, fully contained within the AI region (*η* ∈ [1.5, 3), *g* ∈ [4.5, 6) and *J* ∈ [0.1 *−* 0.25) mV), see blue rectangle in Figure 2. The same amount of training and test data were used as for the full parameter space, so effectively the restricted parameter space was more densely sampled. Estimation errors are shown in Figure 7 (blue lines). With a similarly-sized data set containing only the AI state, the observed error was even smaller than for the full parameter space. Thus focusing on a single network state within which there expectedly is less variation in the LFP, increased the accuracy. However, when using the CNN trained with the data from the *full* parameter space, the estimation accuracy for a restricted test set containing only the AI state, was reduced (Figure 7, purple lines). The accuracy was still better than when estimating parameters across the full parameters space, though, that is, the purple line always was always positioned between the yellow and blue lines in the cumulative plots in Figure 7B. Further, independent of which data set was used, the *g* parameter was always the one with the largest prediction accuracy compared with *η* and *J*.

### 3.3 Highest prediction accuracy of network parameters in AI state

Next, the variation of the parameter estimations errors across the full parameter set was investigated (Figure 8). The estimation of *η* (left panel of Figure 8) was less reliable in the region of low *g* (*g* < 4) which corresponds to the SR state of the network model (Brunel, 2000). The estimation performance of *J* (right panel) was instead worse for the smallest values of *η*, that is, in and around the region of parameters where the network model is in an SI state. The estimation of *g* was generally very accurate for all states of the network (middle panel of Figure 8). Taken together this implies that the highest prediction accuracy of the three network parameters is obtained for the AI state.

**Figure 8:**
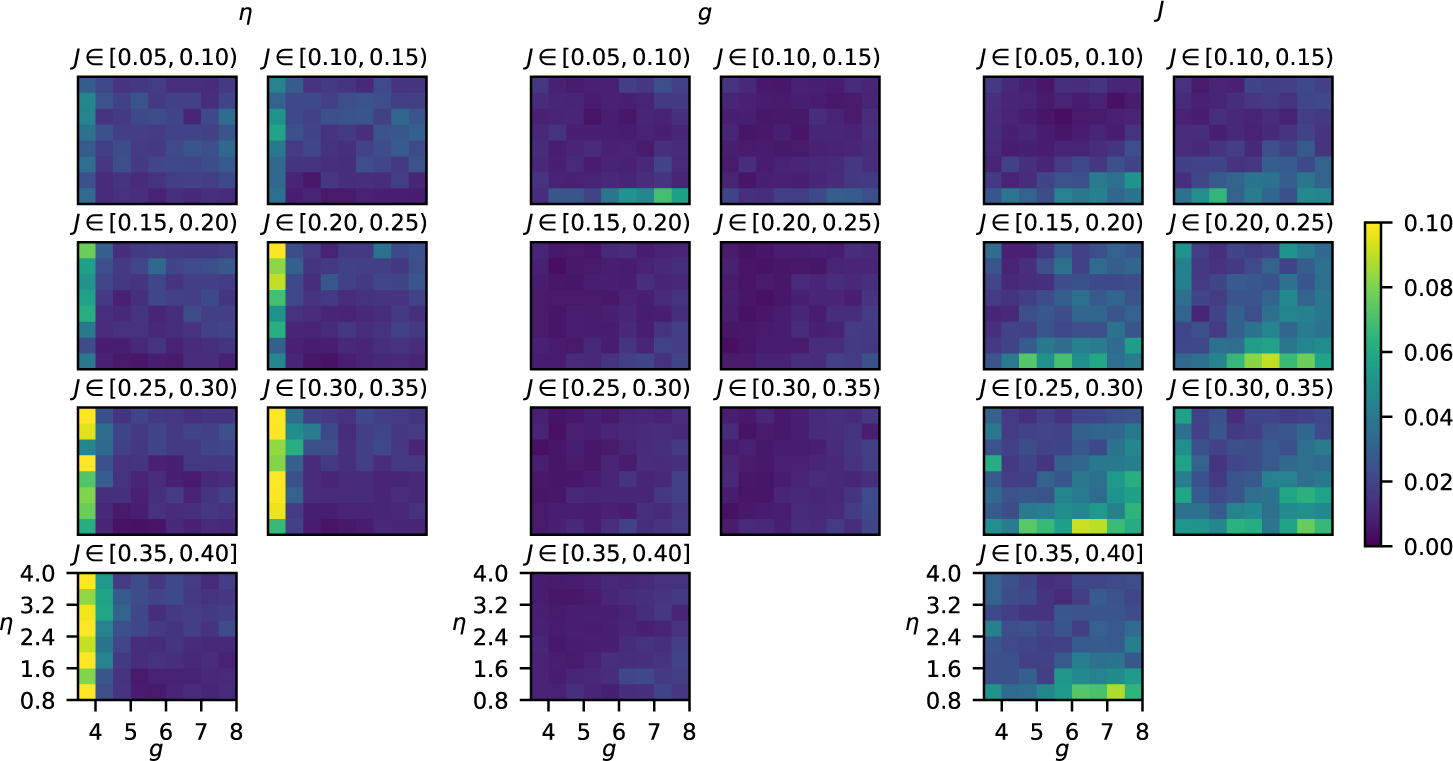
Mean absolute prediction error using full parameter space. Each voxel in the panels shows the error on the validation dataset averaged across the parameter ranges, defined by the pixel size of the grid and the value of *J* indicated above.

We next considered the estimation accuracies across the restricted parameter space corresponding to the AI network state only (*η* ∈ [1.5, 3], *g* ∈ [4.5, 6] and *J* ∈ [0.1 *−* 0.25]), see Figure 9. Also within the AI state, *g* was predicted with the highest accuracy, and *J* had the lowest estimation accuracy. Further, while the estimation accuracy of *g* and *η* was almost constant across the restricted parameter space, the estimation of *J* became worse with increasing values of *J* and *g* (right panel of Figure 9).

**Figure 9:**
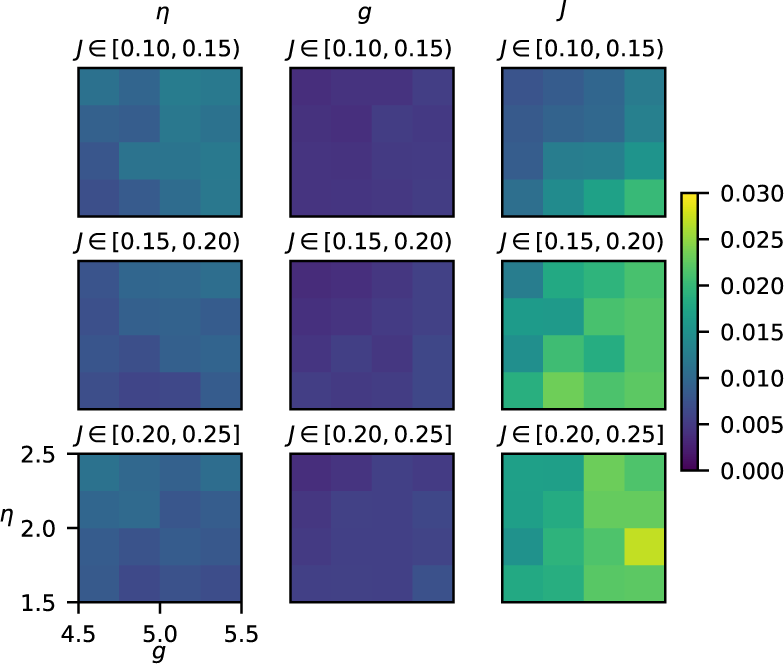
Mean absolute prediction error using restricted parameter set containing only AI state. See caption of Figure 8 for detailed description.

### 3.4 Predicting all parameters at once almost as good as using individually trained CNNs

In the above application all three network parameters were predicted by a single convolutional neural net (CNN). We next investigated to what extent the estimation accuracy changed when CNNs were trained to estimate each parameter separately. The results when considering the full parameter space are shown in Figure 10. As expected the estimation accuracy was always better for these ‘single-prediction’ CNN networks: The error distribution of the *η* prediction was more centered, that is, less biased, for a single prediction network, compared to the ‘combined-prediction’ network (left panel). For the estimation of *g*, the single-prediction network displayed a more narrow peak, also highlighting a slightly better performance. For *J*, the two approaches gave very similar results. Overall, we conclude that merely small gains are achieved for the present application in terms of estimation accuracy by training a separate CNN for each of the three network parameters.

**Figure 10:**
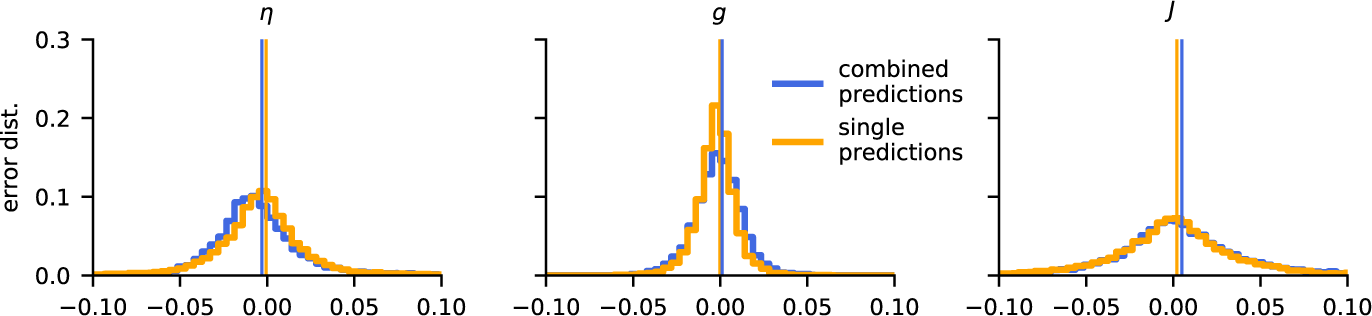
Parameter estimation errors for a single versus multiple CNNs. Comparison of the parameter estimation error, when (i) a single CNN is trained to optimise all three parameters *η, g* and *J* simultaneously (combined predictions), with (ii) three CNNs each trained to estimate a single parameter (single predictions). All parameters were rescaled to the interval [0, 1].

### 3.5 Randomly sampled training-data preferable

The above estimations were based on CNNs trained by LFPs with random network parameters drawn from uniform distributions. To test if the way the parameter space was sampled had an effect on the accuracy of the estimator, we also generated the same amount of training data on a regular grid, spanning the same parameter space and repeated the training. The estimation accuracy was then computed using a randomly generated test data set, and results are shown in Figure 11.

**Figure 11:**
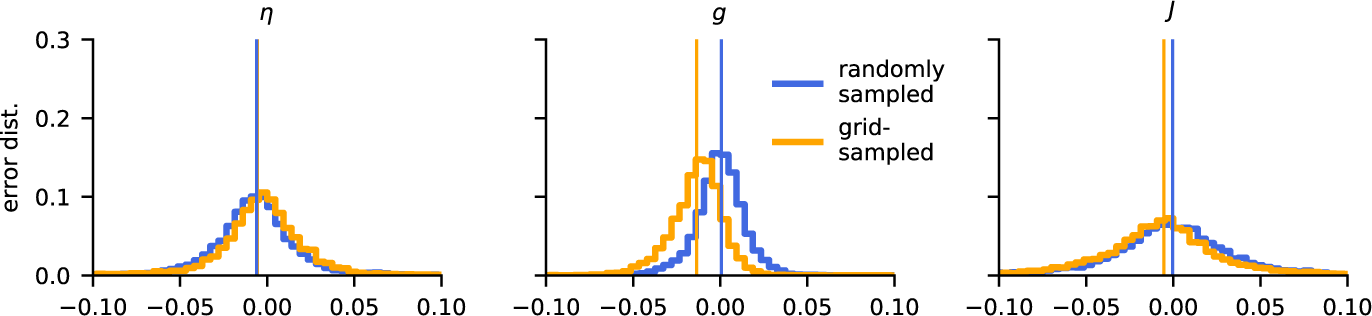
Grid-sampled vs. randomly sampled training data. The plots show error distributions for CNNs trained on data randomly sampled from the parameter set (blue) and from the same amount of training data taken from a regular grid (yellow). All parameters were rescaled to the interval [0, 1].

For the prediction of *η*, there was almost no difference in performance between the CNNs trained with grid-sampled and randomly sampled data (left panel in Figure 11). For *g*, however, the grid-trained data showed a substantial bias towards lower values of *g* (middle panel). Such a bias was also seen in the estimation of *J*, but not so pronounced (right panel).

We speculate that training on grid-sampled data introduces a certain lower resolution to the CNN estimators. Randomly sampled data does not contain such a grid scale and eventually (with sufficient training data) enables the network to learn to interpolate on arbitrary small scales. This intrinsic scale of the grid data might thus be the explanation for the poorer performance of the CNN trained with randomly sampled data.

## 4 Discussion

In the present work we have investigated to what extent the local field potential (LFP), that is, the low-frequency part of an extracellular electrical signal, can be used to extract information about synaptic connection weights and external inputs in the underlying network. As a model we considered the well-known and thoroughly analysed Brunel network comprising an excitatory and an inhibitory population of recurrently connected integrate-and-fire (LIF) neurons (Brunel, 2000). Despite its simplicity, only three parameters (*η*, *g*, *J*) describe external input rate and the weight of the network connections, the model exhibits a high diversity of network dynamics, that is, regular or irregular spiking patterns of individual neurons and synchronous or asynchronous spiking across populations.

The LFP generated by the network was computed using a hybrid scheme (Hagen et al., 2016): Spikes computed by the point-neuron Brunel network where replayed as presynaptic spikes onto biophysically detailed multicompartmental neuron models to compute the LFP as predicted by volume-conductor theory (Lindén et al., 2014; Hagen et al., 2018). We then assessed how well the values of the three model parameters could be estimated from the power spectrum of the stationary ‘background’ LFP signal by application of a convolutional neural net (CNN) (Rawat and Wang, 2017) and indeed found that all parameters could be very accurately estimated. This was the case even when LFPs stemmed from network in different dynamic states (Figures 7,8), but even more so when the LFPs stemmed from the asynchronous irregular (AI) state only (Figure 9).

### 4.1 Generalization to more complex network models

An obvious question is whether the present successful estimation of network parameters from LFPs will extend to more complex network models with more than three parameters specifying the connections like in the Brunel network. Of particular interest here is multilayered cortical network models where several neuronal populations contribute to the LFP signal (Reimann et al., 2013; Głąbska et al., 2014; Tomsett et al., 2015; Głąbska et al., 2016; Hagen et al., 2016).

The estimation problem will expectedly become more difficult as the number of parameters to estimate increases. However, in the present application we only used the power-spectral density (PSD) of the LFP signals from the stationary background state in the parameter estimation. A ‘richer’ LFP signal which may separate the LFP signals for different parameters better, can be obtained by also including the phase information of the LFP Fourier components, but maybe more importantly by also using stimulus-evoked transient LFP signals. Further, in the present application, the parameters were estimated by LFPs from six channels spanning a depth of 0.5 mm. With only a single population contributing to the LFP as in the present case, fewer channels would in fact have sufficed. When several cortical neuronal populations positioned at different depths contribute to the LFP, the spatial variation of the signal contains more information on the network activity. Here the use of a larger number of channels, spanning all cortical layers, should expectedly improve parameter estimation.

To compute the three-second long LFP signals 50000 times to train and test the CNNs in the present study, it was computationally unfeasible to explicitly sum over LFP contributions from each individual presynaptic neuron. Instead we used the approximate formula in Equation 10 based on population firing rates to compute the LFPs, reducing the required computer time by several orders of magnitude. The accuracy of this approximation for the present network was demonstrated for a set of representative examples (Figure 5). In Hagen et al. (2016) where the eight-population Potjans-Diesmann (Potjans and Diesmann, 2014) cortical network model was considered, the same approximation was seen to give fairly accurate LFPs as well (Hagen et al., 2016, Fig. 13), although not as accurate as in the present case as judged by the example tests. Thus the use of the approximation in Equation 10 to compute the LFPs in future applications should be tested on a case-to-case basis.

The choice of using convolutional neural networks (CNNs) within the Keras framework (Chollet et al., 2015) for doing the parameter estimation was made out of convenience. Other machine learning techniques, see Ismail Fawaz et al. (2018) for a recent review, could likely have done as good, or even better. Further, the architecture of the CNNs was not optimised in any systematic way. A systematic study of the best machine learning method to use for LFP-based parameter estimation for more complex network models should be pursued, but is beyond the scope of the present paper.

### 4.2 Implications for analysis of LFPs

For single neurons, biophysics-based modeling is well established (Koch, 1999; Dayan and Abbott, 2001; Sterratt et al., 2011) and numerous biophysically detailed models with anatomically reconstructed dendrites have been made by fitting to experimental data, for example, Migliore et al. (1995); Hay et al. (2011); Halnes et al. (2011); Markram et al. (2015). These models have mainly been fitted to intracellular electrical recordings, but extracellular recordings (Gold et al., 2007) and calcium concentrations (Mäki-Marttunen et al., 2018) can also be used.

Until now the analysis of LFPs have largely been based on statistical methods (Einevoll et al., 2013; Pesaran et al., 2018). An overall goal of the present project is to contribute to the investigation of to what extent LFPs also can be used to develop and validate network models in layered brain structures such as cortex and hippocampus. Spikes have already been used to distinguish candidate network models in cortex (Blomquist et al., 2009; Stimberg et al., 2009), and LFPs recorded *in vitro* have been used to fit hippocampal network models (Chatzikalymniou and Skinner, 2018). There is expectedly a clear link between the accuracy of which a parameter can be (i) estimated from and (ii) fitted to LFP signals. Thus the present observation that network parameters for the Brunel network can be accurately estimated from the background LFPs suggests that the same LFP signal also could be used to accurately fit the same network parameters given that the model structure was known *a priori*. This link between ‘estimatability’ and ‘fitability’ should be properly investigated, not only for the Brunel model, but also for more complex network models. However, such a study is beyond the present scope. A related question that also should be investigated is to what extent LFPs used to distinguish between candidate models with a different network structure, not only different parameters.

### 4.3 Outlook

The recording of single-unit and multi-unit activity (MUA) from the high-frequency part of the extracellular potentials, has historically been the most important method for studying *in vivo* activity in neurons and neural networks. However, the interest in the low-frequency part, the LFP, has seen a resurgence in the last decades. One key reason is the development of new multicontact electrodes allowing for high-density electrical recordings across laminae and areas (as well as computers and hard drives allowing for the storage and analysis of the LFP signals). Another reason is the realisation that the LFP offers a unique window into how the dendrites of neurons integrate synaptic inputs for populations of thousands or more neurons (Lindén et al., 2011). In contrast, the MUA measure the output resulting from this dendritic integration, that is, spikes from a handful of neurons around the electrode contact (Buzsáki, 2004). Thus spikes and LFPs offer complementary information about network activity. Since both signals are produced from the same network model, the combined use of spikes and LFPs appears particularly promising for estimation of network model parameters, or for assessing the merit of candidate network models. Such combined use of spikes and LFPs has been shown to be beneficial in identifying laminar neural populations and their synaptic connectivity patterns from multielectrode cortical recordings (Einevoll et al., 2007; Głąbska et al., 2016). Thus combined use of spikes and LFPs in the estimation of model parameters should be explored in projects where more complex network models are considered where, unlike for the presently considered Brunel network model, the LFP signal is insufficient to alone allow for accurate parameter estimation.

Further, many new optical techniques for probing cortical activity have also been developed and refined, for example, two-photon calcium imaging (Helmchen and Denk, 2005), and voltage-sensitive dye imaging (VSDI), measuring population-averaged membrane potentials (Grinvald and Hildesheim, 2004). Further, at the systems level one has methods such as electroencephalography (EEG) (Nunez and Srinivasan, 2006)), which measures electrical potentials at the scalp, and magnetoencephalography (MEG) (Hämäläinen et al., 1993)) which measures the magnetic field outside the head. These measures can be computed from the activity of candidate network models (Brette and Destexhe, 2012), and tools to facilitate this has been developed (Lindén et al., 2014; Hagen et al., 2018; Gratiy et al., 2018). They can all be used to constrain and validate candidate network models, and used in combination they will likely be particularly powerful.

